# Detection of spatially-localized sounds is robust to saccades and concurrent eye movement-related eardrum oscillations (EMREOs)

**DOI:** 10.1101/2023.04.17.537161

**Authors:** Felix Bröhl, Christoph Kayser

## Abstract

Hearing is an active process and recent studies show that even the ear is affected by cognitive states or motor actions. One example are movements of the eardrum induced by saccadic eye movements - known as “eye movement-related eardrum oscillations” (EMREOs). While these are systematically shaped by the direction and size of saccades, the consequences of saccadic eye movements and their resulting EMREOs for hearing remain unclear. We here studied their implications for the detection of near-threshold clicks in human participants. Across three experiments sound detection was not affected by their time of presentation relative to saccade onset, by saccade amplitude or direction. While the EMREOs were shaped by the direction and amplitude of the saccadic movement, inducing covert shifts in spatial attention did not affect the EMREO, suggesting that this signature of active sensing is restricted to overt changes in visual focus. Importantly, in our experiments fluctuations in the EMREO amplitude were not related to detection performance, at least when monaural cues are sufficient. Hence while eye movements may shape the transduction of acoustic information the behavioral implications remain unclear.

## 1. Introduction

To perceive and navigate our environment the brain needs to coordinate acoustic and visual spatial signals in order to bind percepts arising from a common object. This poses an intricate challenge as the eyes move relative to the head and ears, displacing the spatial reference frames for hearing and vision (Groh and Sparks, 1992; Kopinska and Harris, 2003; Mullette-Gillman et al., 2005; Krüger et al., 2016). One way by which our brain may solve this challenge is to engage the auditory system in active perception, for example by modulating sound encoding in relation to eye or body movements.

Recent work has shown that this process may even influence the ear itself. A pioneering study by Gruters and colleagues showed that our eardrums transiently vibrate following the execution of saccadic eye movements - a phenomenon termed ‘eye movement related eardrum oscillations’ or EMREO (Gruters et al., 2018). These EMREOs exhibit functional specificity as they are systematically related to the direction and amplitude of saccades. More so, these EMREOs have been shown to reliably contain information about initial fixation points and saccade direction in two-dimensional space (Murphy et al., 2020; Lovich et al., 2022). Incoming sounds are transduced by setting the eardrum in vibration, therefore one may expect that these EMREOs shape the transduction of sounds and thereby how they are perceived. Because EMREOs in both ears appear to vibrate simultaneously in opposing phases, it has been speculated that EMREOs may affect interaural cues and could selectively modulate perception depending on the congruency between saccade direction and sound location (Gruters et al., 2018; Cho et al., 2023). Furthermore, hearing entails not only the reflexive encoding of incoming sounds but also the expectation of those. Hence processes related to spatially guided attention may possibly influence the auditory periphery as well (Köhler et al., 2021; Gehmacher et al., 2022), and a modulation of EMREOs could be one possible implementation.

An influence of eye movements on the auditory pathways has been observed at multiple stages, from the inferior colliculus to the auditory cortex (Groh et al., 2001; Werner-Reiss et al., 2003; Fu et al., 2004; Maier and Groh, 2010; Lee and Groh, 2012; Barczak et al., 2022). Yet studies on the relation between oculomotor behavior and auditory perception have yielded conflicting evidence. While some suggest a relation between fixation position and systematic errors in localizing sounds (Lewald, 1997, 1998; Lewald and Ehrenstein, 1998), others suggest that the presence of a visual target in itself can contribute to such errors (Klingenhoefer and Bremmer, 2009) as dissecting potential influences of a visual stimulus itself from oculomotor behavior can be difficult (Harris and Lieberman, 1996). Furthermore, when presenting targets during a saccade, localization of visual targets was similarly erroneous as for acoustic targets (Collins et al., 2010; Krüger et al., 2016). Hence, despite the presence of oculomotor signals along the auditory pathway the influence of oculomotor behavior on hearing remains unclear.

We directly tested the relation between saccadic eye movements, concurrent EMREOs and perception across three experiments in human participants. In a first experiment we replicated the existence of EMREOs and confirmed their directional selectivity. In a second experiment we probed how the detection of near-threshold sounds is shaped when presented before, during or after a saccade, and whether detection performance is modulated by the amplitude of the EMREO signal. Finally, in a third experiment we used spatial cueing to probe whether EMREOs are shaped by covert changes in spatial attention and whether this affects task performance via the modulation of EMREOs. Our data confirm that EMREOs are evoked in a stereotypical manner by saccadic eye movements, but do not offer evidence that saccades or their induced EMREOs modulate the detection of spatially localized sounds. Furthermore, we found that the EMREO time course was robust towards manipulations of spatial attention. All in all our results suggest that tone detection remains unaffected by saccadic eye movements and their induced EMREOs.

## 2. Materials and Methods

### Participants and data acquisition

We collected data from 26 volunteers recruited among university students (18 female, 8 male, 0 diverse, mean age = 25.6 ± 3.8 SD). All participants were screened using the quick hearing questionnaire (Kochkin and Bentler, 2010) and provided written informed consent. During the study 14 participants were excluded subsequent to experiment 1 as detailed below. The study was approved by the local ethics committee of Bielefeld University, Germany, and was conducted in accordance with the Declaration of Helsinki. The whole study took about three hours and participants received monetary compensation of 10 € per hour.

Participants were seated in a dimly lit and acoustically shielded room and 100 cm in front of an acoustically transparent screen (Screen International Modigliani, 2×1 m) with their chin on a chin rest. Speakers (Monacor MKS-26/SW, MONACOR International GmbH, Germany) were located behind the screen at -6, and 6 degrees azimuth. Sounds played from these speakers were generated using a Creative soundblasterZ Soundcard (sampling rate 44,100 Hz) and amplified using a t.amp E4-130 amplifier (Thomann Germany). An additional route of sound presentation was implemented via the DPOAE amplifier (see below). Here sounds were generated using the same sound card. Visual stimuli were projected onto the screen (Acer Predator Z650, Acer Inc., Taiwan) and stimulus presentation was controlled using the Psychophysics toolbox (Brainard, 1997) for MATLAB (The MathWorks Inc., Natick, MA). The accuracy of timing was manually tested using an oscilloscope.

Ear data were acquired using Etymotic 10B+ in-ear microphones (Etymotic research) with foam ear buds. The appropriate earbud size was determined by visual inspection. Earbuds and microphones were fit into both ears and held into place by skin-compatible sticky tape, though data was only recorded from the right ear in all participants. The microphone signal was preamplified using a DPOAE amplifier (ER-10C; Etymotic; with gain set to +20 dB), digitized using an ActiveTwo AD-box (BioSemi) and stored at a sampling rate of 16,384 Hz. Participants’ eye movements were continuously recorded from their left eye with an EyeLink 1000 Plus eye tracking system (SR research) at 1000 Hz (using the EyeLink 1000 “cognitive” setting). The eye tracking system was calibrated before each experiment and block using a 5-point procedure.

### Experimental design

The study consisted of five phases: (1) Click-evoked otoacoustic emission, (2) Determination of sound detection thresholds, (3) experiment one, (4) experiment two and (5) experiment three.

#### Click-evoked otoacoustic emissions

To assert whether the microphones were fit correctly and to screen for an impairment of outer hair cell function, we collected click-evoked otoacoustic emissions by presenting 100 consecutive 0.1 ms positive monophasic bursts via the in-ear microphone (Probst et al., 1991). Each two click tones were presented at a semi-random interval of 400 ms fixed duration plus an additional interval (0 to 200 ms, uniformly sampled). An additional 15 s of baseline silence was recorded. Participants were instructed to move as little as possible and to fixate a crosshair at the center of the screen. The subsequent visual inspection showed that all participants featured robust otoacoustic emissions.

#### Determination of sound detection threshold

We determined the detection threshold for the task used in experiments 2 and 3. Participants were asked to detect target sounds (1 ms positive monophasic bursts, ‘clicks’) of variable intensity that were presented from the speakers behind the projector screen. For each participant we determined the detection threshold using three interleaved 2-down-1-up staircases (stopping after 10 reversals). The average of these three thresholds was used for the main experiments.

##### Experiment 1

The goal of this experiment was to replicate the emergence of EMREOs in the absence of an auditory task. Following previous work (Gruters et al., 2018), participants were asked to fixate on a small white dot of 0.2 degrees radius, whose position jumped along the vertical and horizontal axes on the screen. At the start of each trial the white dot appeared at the center of the screen and after 500 ms of continued fixation moved to either -12, -6, 6, or 12 degrees up, down, left or rightwards (Fig. 1). The positions at [-12, -6] degrees are contralateral to, those at [6, 12] degrees are ipsilateral to the recorded right ear. In the successive trial the dot moved from the eccentric position back to the center of the screen. Although we recorded both vertical and horizontal saccades that moved outwards and inwards in this experiment, we focused only on outward horizontal saccades for the main analysis and subsequent experiments. Inter-trial intervals were 900 ms plus a random period (0 to 500 ms; uniformly sampled). This experiment comprised four blocks of 160 trials.

**Fig. 1.**
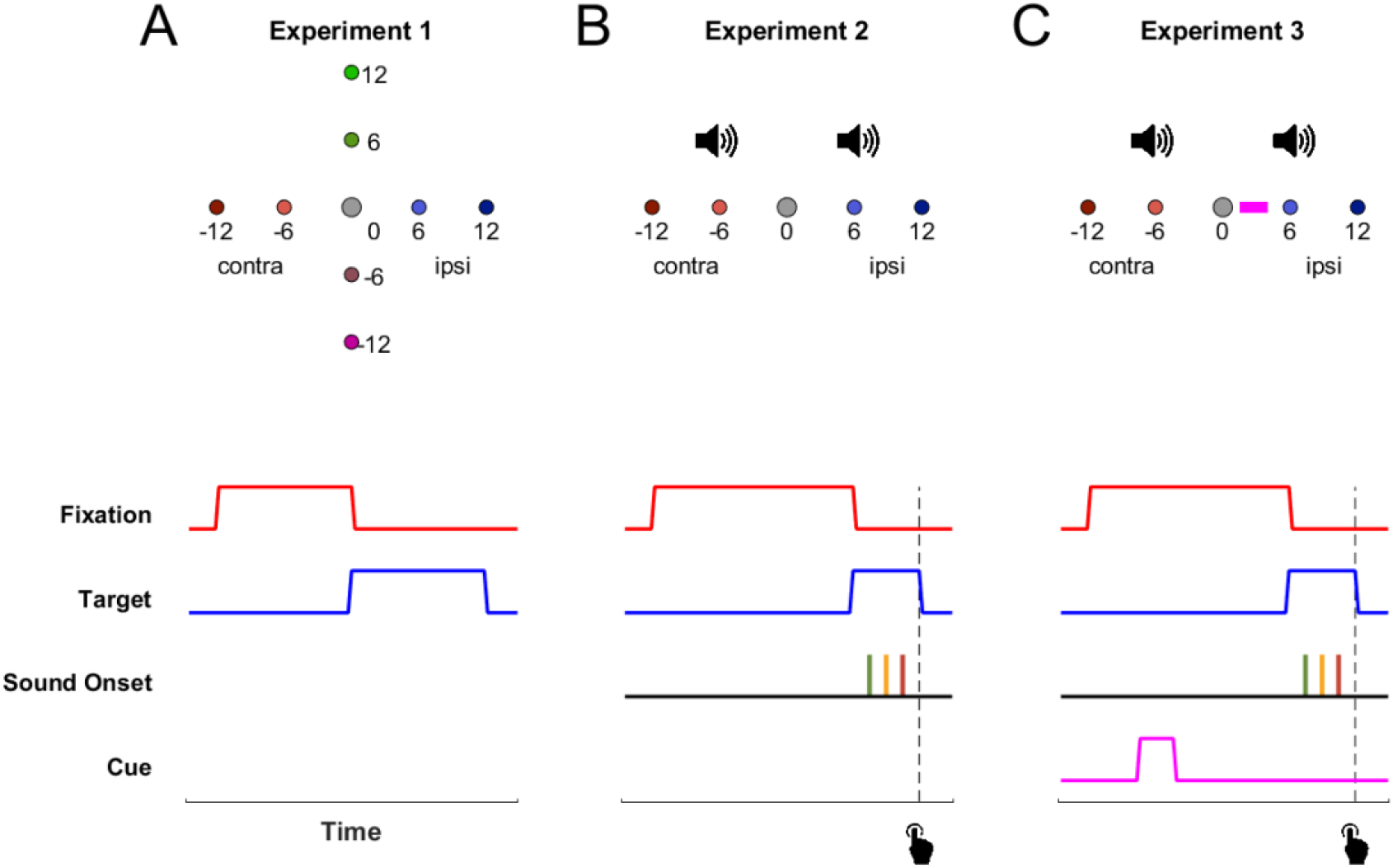
Experiment setup and trial structure. The diagrams depict the spatial (top) and temporal (bottom) trial structure for each experiments. (**A**) In experiment 1, participants were presented with a dot at the center of the screen, which randomly moved to a new location at [-12, -6, 6, 12] degrees either vertically or horizontally from the central fixation point. In the subsequent trial the dot moved back to the center. (**B**) In experiment 2, targets were located on the horizontal line to the left or right of the central fixation point. During each trial the dot moved to a new randomly chosen eccentric position while a sound was presented from either -6 or 6 degrees in 70% of trials. The participant’s task was to detect a near threshold click sound. This click was played either before (green), during (yellow) or after (red) of the saccade. (**C**) Experiment 3 in addition featured a spatial cue that predicted the correct sound location on 70% of the trials. The cue was a bar (depicted here in pink) flashed for 250 ms next to the central fixation point. In all experiments the occurrence of each eccentric visual target was equally likely. Positions at [-12, -6] degrees are contralateral to the recorded right ear.

Based on the outcome of this experiment we excluded participants from the remaining study if either the quality of eye tracking or the microphone signal were not sufficient, which was based on visual inspection of the continuous signal power during the recording. Practically, we excluded 12 participants. From the eye tracking data obtained in this experiment we also determined the participant-wise mean saccade onset latency (P_SOL_) and mean saccade duration (P_SD_) which were used to define the sound onsets times for experiments 2 and 3.

##### Experiment 2

This experiment was designed to probe sound detection relative to eye movements and EMREOs. Participants executed visually guided horizontal saccades similar to experiment 1 and performed a sound detection task. A white dot appeared at the center of the screen and after fixation period (1250 ms plus a uniform random period of up to 500 ms) the dot moved to another location (at -12, -6, 6, or 12 degrees azimuth). In 70% of trials a click tone was presented from either the speaker at -6 or 6 degrees and at one of three time points: before, during or after the saccade. The timing of the sound onset was based on the saccade latencies and durations for each individual participant (P_SOL_ and P_SD_) obtained from experiment 1 and was defined as follows:

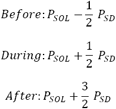

Note that this does not result in sounds being strictly presented in the respective saccade time bin, since the sounds were presented at a fixed participant dependent latency, but saccade onsets naturally occurred with varying latencies across trials. We used this mode of presentation to achieve a roughly comparable amount of sounds played in either time bin. For analysis we grouped the trials according to the actual sound to saccade latency. Both the sound position and the sound omittance were counterbalanced across saccade times and directions. Participants were asked to follow the dot with their eyes and to indicate the detection of the sound via a button press as quickly as possible. Trials with no responses within 2 s after sound presentation were counted as a trial where no sound was detected. Because the foam earbuds may slightly deform during the experiment and could change detection thresholds we updated the click intensity every ten trials to either stay constant or change by ± 5 % of its prior value to assert that the detection accuracy does not decrease below 55 % or increase above 75 %. This experiment consisted of four blocks of 144 trials.

##### Experiment 3

The third experiment probed whether EMREOs and sound detection are shaped by covert spatial attention. The experiment was structured similarly to experiment 2 with the addition of a spatial cue. This cue predicted the upcoming sound location correctly in 70% of trials, termed a valid cue, and did so incorrectly in 30% of trials, termed an invalid cue. At the start of each trial participants had to fixate a white dot in the center of the screen. After 50 ms the spatial cue was presented in the form of a white bar of 2.5 degrees length and 0.2 degrees width flashed for 250 ms to the left or to the right of the dot. Subsequently, the dot remained at the central position (900 ms plus a random period of 0 to 500 ms, uniformly sampled) until it moved to one of the eccentric locations (-12, -6, 6, or 12 degrees). Participants were instructed to keep fixation on the dot while the cue was presented and were told that it was presented in each trial, whether a sound was played or not. This experiment consisted of four blocks of 144 trials.

### Data preprocessing and analysis

#### Preprocessing

The microphone and eye tracking data were resampled to a common sample rate of 2 kHz. The microphone data was then aligned to the saccade onsets reported by the eye-tracking system (using the EyeLink 1000 “cognitive” setting, velocity threshold: 30°/s, acceleration threshold: 8000°/s). Outlier trials were removed based on the following criteria pertaining to the quality of eye tracking and the microphone data: (1) the standard deviation of the microphone signal over the trial exceeded 2.5 SD of the signal obtained across all trials, (2) the saccade onset time exceeded 500 ms, (3) the initial fixation at the start of the saccade was outside 3 SD from the fixated position computed across all trials, (4) during any time of the trial the eye position moved outside of a 16 degrees azimuth boundary around the center point (which indicate blinks or otherwise faulty eye tracking). Across participants 91.77 ± 8.94 % of trials were included. These trials were grouped by saccade direction and amplitude ([-12, -6, 6, 12] degrees). Within experiment 1 participants also executed saccades in the vertical direction, which we analyzed separately, and performed saccades towards the center, which were excluded from further analysis.

### Regression of EMREOs against eye movement parameters

To quantify the dependency of the EMREO signal on eye movement parameters we implemented a linear model predicting the EMREO amplitude at each time point based on the horizontal eye displacement and a constant factor. As previous work has indicated that the individual saccade start and endpoints may also contribute to the EMREO (Murphy et al., 2020), we added two further predictors: the variance of horizontal eye position at the start of the saccades and the trial-wise deviation from the visual target. The model can then be described as

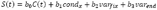

where *S*(*t*) denotes the microphone signal and *C*(*t*) the constant term over time, *cond*_*x*_ the conditional horizontal eye displacement and *var*_*fix*_ and *var*_*end*_ the horizontal eye position variance at the start and end of the saccade, respectively. The latter was derived by subtracting the target location from the actual saccade endpoint. To determine the unique variance explained by each predictor we computed partial regression models that left out individual predictors. The variance attributed to each predictor was determined by the difference between the full model and the reduced model.

### Analysis of behavioral data

Trials were split by saccade direction (contra-/ipsilateral), amplitude (±6/±12 degrees), sound location (contra-/ipsilateral) and by saccade-sound congruency (congruent/incongruent; defined by whether the saccade was moving towards the same side as the sound was played). For each constellation we characterized behavioral performance using sensitivity (d prime) and bias. We also computed the median reaction time for correct trials. For experiment 3 we also separated trials by the validity of the spatial cue (valid/invalid) and the cue-saccade congruency (congruent/incongruent), which indicates whether the cue and the saccade were guided in the same direction. In a separate analysis, we compared sensitivity and bias between trials in which the target sound was presented before, during or after the saccade onset. For reaction times we computed a linear regression of the log-transformed reaction times against the temporal delay between saccade onset and the sound presentation time. Finally, to directly link EMREO amplitude and detection performance we focused on trials in which the sound was presented (or omitted) during a saccade. These were partitioned by response (correct/incorrect) and saccade condition ([-12, -6, 6, 12] degrees). To reduce the EMREO time course to a single amplitude measure we computed the power of the average EMREOs. This was done by taking the absolute of the Hilbert transform of the participant and condition-averaged time course and subsequently averaged the EMREO power within the 0 to 75 ms window.

### Statistical analysis

For regression models the significance of individual predictors was tested using a cluster-based permutation procedure (Nichols and Holmes, 2003). As the overall time course of the EMREOs diminishes around 75 ms after saccade onset, we focused our analyses on the time window from 0 to 75 ms. We first compared individual predictors at each time point against zero using a two-sided student’s t-test across participants. The outcome was thresholded at a level of p<0.05 and clusters of significant time points (of at least 2 ms duration) were compared to randomization distribution obtained by flipping the sign of the t-test randomly 10,000 times. The same approach was applied to compare the EMREOs between conditions based saccade targets and response labels, as well as cue validity and congruency. The effect size for significant clusters were computed from the peak test statistic. To test the relation between the overall EMREO amplitude and factors such as saccade direction we relied on a ANOVAs. To compare sound detection sensitivity, detection bias and median reaction times between conditions across participants we relied on two-sided paired t-tests against zero. The p-values of these were adjusted for multiple tests using the Benjamini-Hochberg procedure (Benjamini and Hochberg, 1995). To compare correct detections among the three bins of sound onsets (before, during and after saccade) we relied on a one-way repeated measures ANOVA. The regression weights of the reaction time to the sound onset relative to the saccade onset was tested using a two-sided paired t-test against zero. Again the resulting p-values were corrected for multiple testing using the Benjamini-Hochberg procedure. For all statistical tests we provide exact p-values, test statistics and measures of effect size. For all parametric tests we transformed test-statistics into Bayes Factors (BF) as BF_10_ by using BIC approximation (Krekelberg, 2022). For easier interpretation we transformed all BF_10_ values below 1 into -BF_01_.

## 3. Results

### EMREOs are shaped by saccade target rather than saccade variability

Participants performed visually guided saccades along the horizontal or vertical axes while we recorded eardrum activity using an in-ear microphone from their right ears. To visualize the EMREO we aligned the microphone signal to saccade onsets and obtained the trial-averaged signal for each saccade direction and amplitude. Figure 2 shows the resulting EMREOs for individual participants (panel **A**) and as group average (panel **B**). For horizontal saccades, these reveal the expected and consistent patterns across participants, including opposing phases for the two saccade directions and dependency of EMREO amplitude on saccade amplitude. The latencies of the first two peaks of the EMREOs occurred after 14.9 ± 3.4 ms and 21.0 ± 7.9 ms (mean ± SD). The vertical saccades (performed only in experiment 1) reveal a similar albeit less clear relationship between EMREO phase and saccade direction, as expected based on previous reports (Gruters et al., 2018; Murphy et al., 2020; Lovich et al., 2022).

**Fig. 2.**
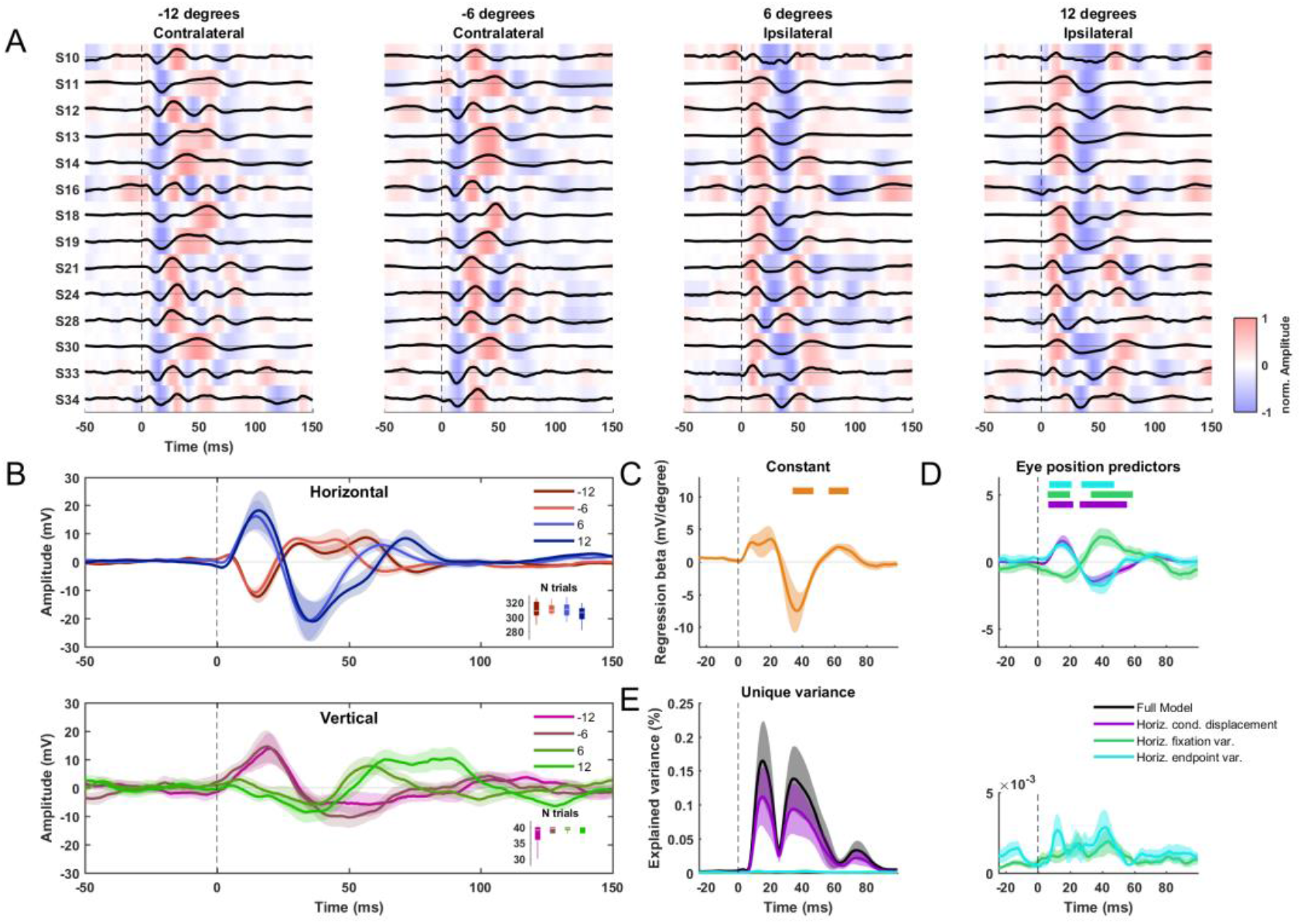
EMREOs contain information about saccade targets. (**A**) EMREO time course (obtained from the right ears) for individual participants and the four horizontal conditions aligned to saccade onsets (dashed vertical lines). Saccade positions at [-12, -6] are contralateral and positions at [6, 12] are ipsilateral to the recorded ear. Black lines indicate the average across all three experiments, normalized to one within participants. (**B**) The grand average EMREO trace across all participants (mean and SEM across participants) for horizontal (top) and vertical (bottom) saccades. The scaling of EMREO amplitude with saccade size is best visible for horizontal saccade and the EMREOs’ first inflection (within 25 ms after saccade onset). The small inlays indicate the average number of trials included per condition across participants. (**C**,**D**) Weights for the constant (**C**) and each predictor based on eye tracking data (**D**) that were used to predict the EMREOs time course across conditions. Thick bars indicate clusters of time points significantly different from zero (cluster-based permutation procedure). (**E**) Unique variance explained by individual predictors. The right panel visualizes horizontal fixation and endpoint variance on a smaller scale for better comparison. Lines show the group average, shaded areas the standard error of the mean across participants.

The scaling of the EMREO with saccade direction and amplitude could pertain to the instructed saccade vector that was indicated on the screen, or could directly pertain to the actually executed movement. The latter may differ from the visual instruction by a difference in the precise starting position (within the allowed tolerance) and an over-or under-shoot at the landing position. We probed which of these parameters hold significant predictive power for the EMREO time course using linear modelling. This model included the conditional horizontal eye displacement and the horizontal variability around the initial fixation point (0.83 degrees SD) and around the targets ([1.76, 1.41, 1.17, 1.86] degrees SD for the -12 to 12 degrees conditions) as predictors. This revealed a significant cluster for the conditional displacement (Fig. 2, **D**, 6.5 to 22.0 ms, p=0.02 maxsum=114.36 cohen’s D_peak_=1.00; and 26.0 to 55.5 ms, p=0.001 maxsum=-192.31 cohen’s D_peak_=0.68), the starting variability (Fig. 2, **D**, 6.0 to 20.0 ms, p=0.04 maxsum=-88.64 cohen’s D_peak_=1.00; 33.0 to 59.0 ms, p=0.004 maxsum=141.14 cohen’s D_peak_=0.63) and the endpoint variability (Fig. 2, **D**, 7.0 to 21.0 ms, p=0.039 maxsum=94.56 cohen’s D_peak_=1.00; and 27.0 to 47.5 ms, p=0.011 maxsum=-117.43 cohen’s D_peak_=0.68). The constant, reflecting a consistent EMREO component independent of saccade direction, featured two significant clusters (**C**, 34.0 to 47.0 ms, p=0.035 maxsum=-83.94 cohen’s D_peak_=0.60; and 56.5 to 69.0 ms, p=0.044 maxsum=76.35 cohen’s D_peak_=0.22). Comparing the amount of variance in the EMREO signal explained by each predictor revealed that the conditional eye displacement explained by far the most, while the precise start and end-points of the eye movement contribute consistently but with much smaller explained variance to the EMREO (Fig. 2 **E**).

### Sound detection performance is not affected by saccadic eye movements

In experiments 2 and 3 we asked participants to detect a near-threshold click that was presented before, during or after a saccade. We quantified participants’ performance using detection sensitivity and bias and their response times. Across these measures we found that participants’ behavior was not affected by the direction and amplitude of the saccade or the location of the sound in either experiment (Fig. 3, Tab. 1).

**Fig. 3.**
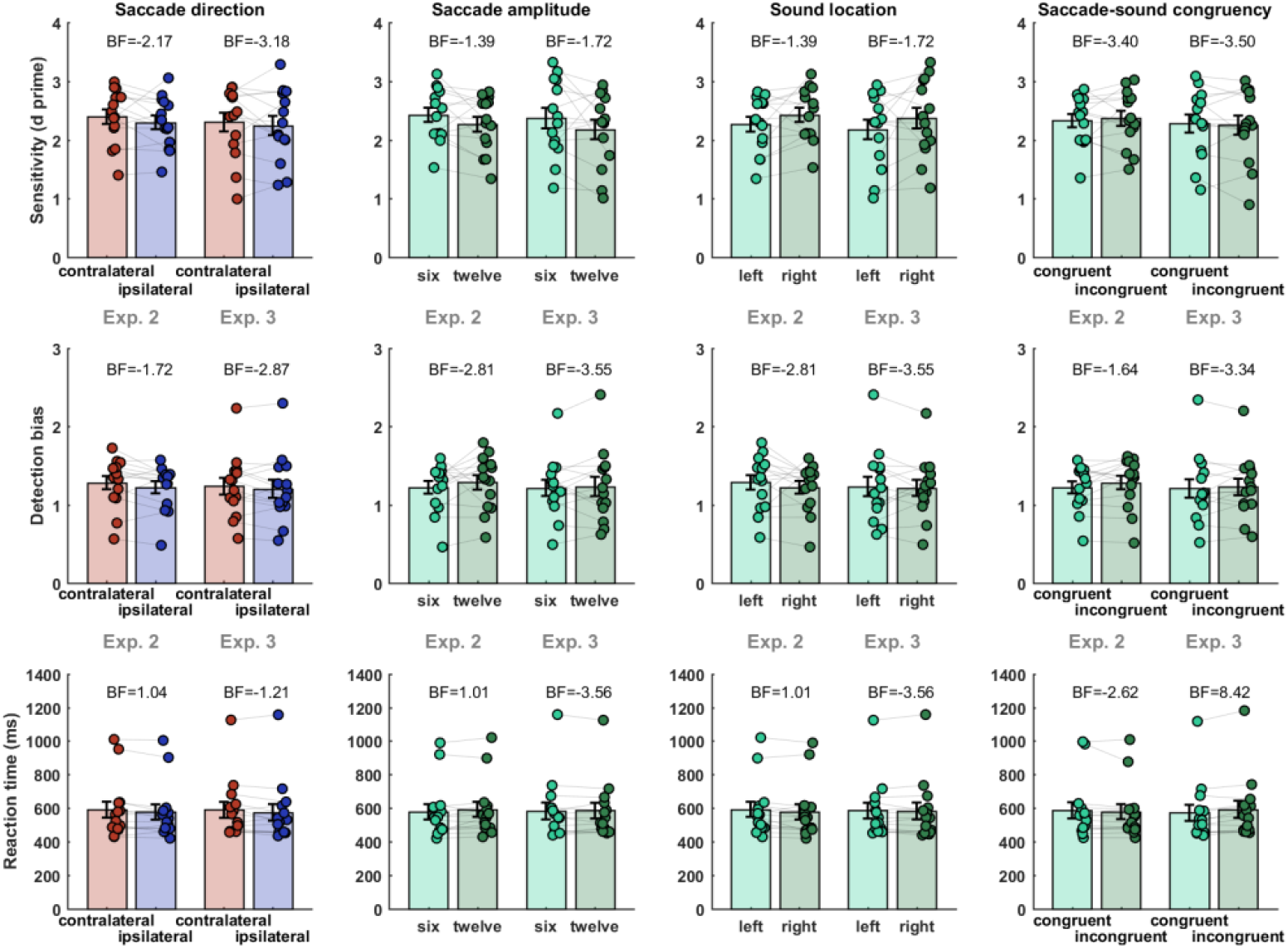
Sound detection is not affected by saccade parameters. Detection sensitivity (top row), bias (middle row) and reaction times (bottom row) were determined for trials split by saccade direction, saccade amplitude, sound location and saccade-sound congruency (i.e., whether the saccade was directed at the same direction the sound was presented from). In each panel, the left two bars indicate data from experiment two, the right two bars indicate data from experiment three. Dots represent individual participants. Statistical comparisons within experiments and factors were based on two-sided paired t-tests for which we report Bayes factors.

**Table 1.** Statistical results for sound detection. For each saccade parameter we tested its effect on sensitivity, bias and reaction times across participants using a two-sided paired t-test. T-statistics were transformed into Bayes factors (BF) and the p-values were subsequently FDR-corrected within this table.

|  |  | Saccade direction |  | Saccade amplitude |  | Sound location |  | Saccade-sound congruency |  |
| --- | --- | --- | --- | --- | --- | --- | --- | --- | --- |
|  |  | Exp.2 | Exp. 3 | Exp.2 | Exp. 3 | Exp.2 | Exp. 3 | Exp.2 | Exp. 3 |
| Sensitivity | $p_{FDR}$ | 0.56 | 0.77 | 0.43 | 0.43 | 0.43 | 0.43 | 0.77 | 0.77 |
|  | BF | -2.17 | -3.18 | -1.39 | -1.72 | -1.39 | -1.72 | -3.40 | -3.50 |
|  | Cohen's D | 0.30 | 0.16 | 0.41 | 0.36 | 0.41 | 0.36 | 0.12 | 0.10 |
| Bias | $p_{FDR}$ | 0.43 | 0.69 | 0.69 | 0.77 | 0.69 | 0.77 | 0.43 | 0.77 |
|  | BF | -1.72 | -2.87 | -2.81 | -3.55 | -2.81 | -3.55 | -1.64 | -3.34 |
|  | Cohen's D | 0.36 | 0.20 | 0.21 | 0.08 | 0.21 | 0.08 | 0.38 | 0.13 |
| RT | $p_{FDR}$ | 0.43 | 0.43 | 0.43 | 0.77 | 0.43 | 0.77 | 0.69 | 0.15 |
|  | BF | 1.04 | -1.21 | 1.01 | -3.56 | 1.01 | -3.56 | -2.62 | 8.42 |
|  | Cohen's D | 0.50 | 0.45 | 0.49 | 0.08 | 0.49 | 0.08 | 0.24 | 0.87 |

In experiment 3 differences in reaction times were faster when the saccade was directed towards the side where the target sound was presented (Fig. 3, saccade-sound congruency: T(13)=3.26, p_FDR_=0.1, BF=8.42, cohen’s D=0.87). This likely is the result of the presence of the informative cue in that experiment.

The experiments were designed such that sounds were played either before, during or after the saccade. For the analysis we grouped trials based on the actual relative timing of sound presentation. Comparing sensitivity between these three epochs revealed evidence against an effect of epoch for both experiments (Fig. 4 **B**, Exp. 2, F(2,39)=0.22, p_FDR_=0.96, BF=-33.18; **C**, Exp. 3, F(2,39)=0.42, p_FDR_=0.96, BF=-26.94) and the same was found for bias (Exp. 2, F(2,39)=0.71, p_FDR_=0.96, BF=-12.94; Exp. 3, F(2,39)=0.15, p_FDR_=0.96, BF=-12.07). To probe an effect of sound-saccade timing for reaction times we used a linear regression of reaction times against the sound onset time. This revealed no effect for experiment two (Fig. 4 **B**, T(13)=-0.48, p_FDR_=0.89, BF=-3.35, cohen’s D=0.13). In experiment three, participants responded faster when the sound was presented later in the trial (Fig. 4 **C**, T(13)=-4.59, p_FDR_=0.002, BF=70.51, cohen’s D=1.23), which we again interpret as an effect of the cue in this experiment. All in all, this data hence suggests that the timing of the sound relative to saccadic eye movements has no consistent influence on sound detection.

**Fig. 4.**
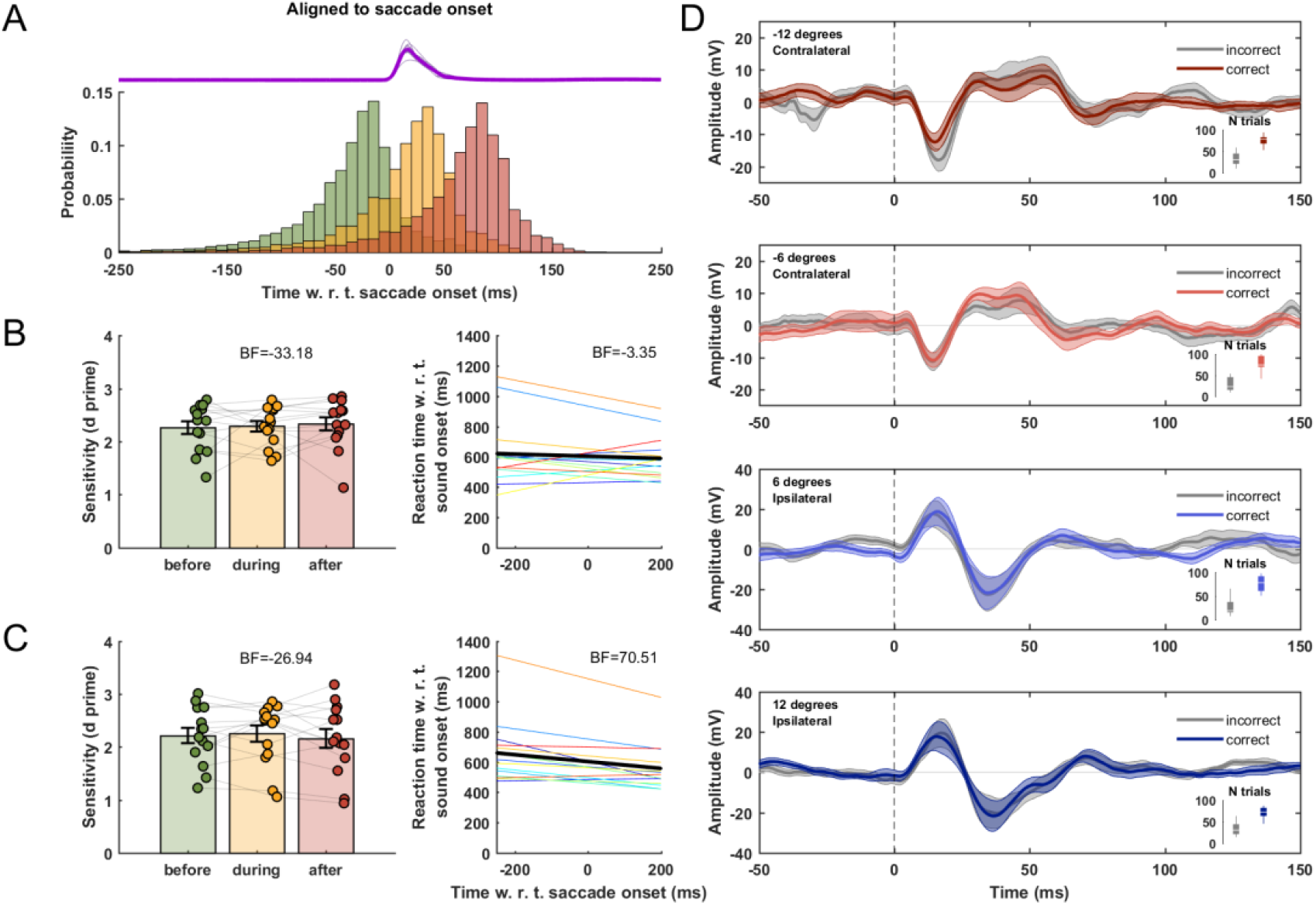
Relationship between EMREOs and sound detection. (**A**) Distribution of sound presentation times relative to saccade onsets across participants. Magenta traces on top indicate the mean eye velocity per participant, the thick magenta line indicates the group average. (**B**,**C**) Left: Sensitivity for each time bin in experiment two (**B**) and experiment three (**C**). Sensitivity was compared across time bins using an ANOVA. Right: regression weights of reaction times against the time delay between sound presentation and saccade onset. Individual lines represent individual participants and the thick black line indicates the mean across participants. Regression weights were tested for significance using a two-sided t-test. (**D**) EMREOs split by saccade target condition ([-12, -6, 6, 12] degrees) and separately for trials with correct and wrong responses. This analysis includes only trials where a sound was presented during the saccade and combined data from experiments 2 and 3. Lines indicate the mean EMREO time course across participants for correct (colored) and incorrect (gray) trials, shaded areas indicate the standard error of the mean.

### Sound detection is not related to EMREO fluctuations

To probe a direct relation between the EMREO amplitude and behavior we compared EMREOs in experiments 2 and 3 combined between trials featuring correct and wrong responses. For this analysis we focused on trials where a sound occurred (or was omitted) during the saccade and separated these based-on saccade target ([-12, -6, 6, 12] degrees) and response (correct/incorrect). For a statistical analysis we computed the average EMREO power and contrasted this between conditions and responses using a repeated-measures ANOVA. This revealed a significant effect of saccade condition on the EMREO, although there was modest evidence for the alternative of no difference (Fig. 4, **D**, F(3,104)=4.072, p=0.009, BF=-2.36). In addition, this revealed no effect of responses (correct vs. wrong; F(1,104)=1.2, p=0.28, BF=-5.57) nor an interaction between correctness and saccade condition (F(3,104)=0.3, p=0.82, BF=-728.29).

### EMREOS are robust to covert spatial attention

In the third experiment we asked whether spatial attention shapes the subsequent EMREO. To test this we provided a spatial cue that correctly predicted the direction of the upcoming sound in 70 % of the trials. We then analyzed the data based on cue validity (whether the sound was actually presented in the cued direction), and cue-saccade congruency (whether the cue and saccade point in the same direction). We found no evidence that cue validity shaped the EMREO, nor that the congruency between cue and saccade direction modulated the EMREO time course (Fig. 5 **A** and **B**, left panels). To directly compare the EMREO amplitude between conditions, we computed the mean EMREO power and tested the influence of cue-sound validity and cue-saccade congruency. This revealed, as expected, an effect of saccade direction (Fig. 5, **A**, F(1,52)=6.76, p=0.012, BF=4.09; **B**, F(1,52)=5.46, p=0.023, BF=2.19), but no effects of cue validity (**A**, F(1,52)=0.0026, p=0.96, BF=-7.47) or cue-saccade congruency (**B**, F(1,52)=0.044, p=0.83, BF=-7.31) nor an interaction between saccade direction and cue-sound validity (**A**, F(1,52)=0.012, p=0.91, BF=-7.43) or saccade direction and cue-saccade congruency (**B**, F(1,52)=0.0003, p=0.99, BF=-7.48).

**Fig. 5.**
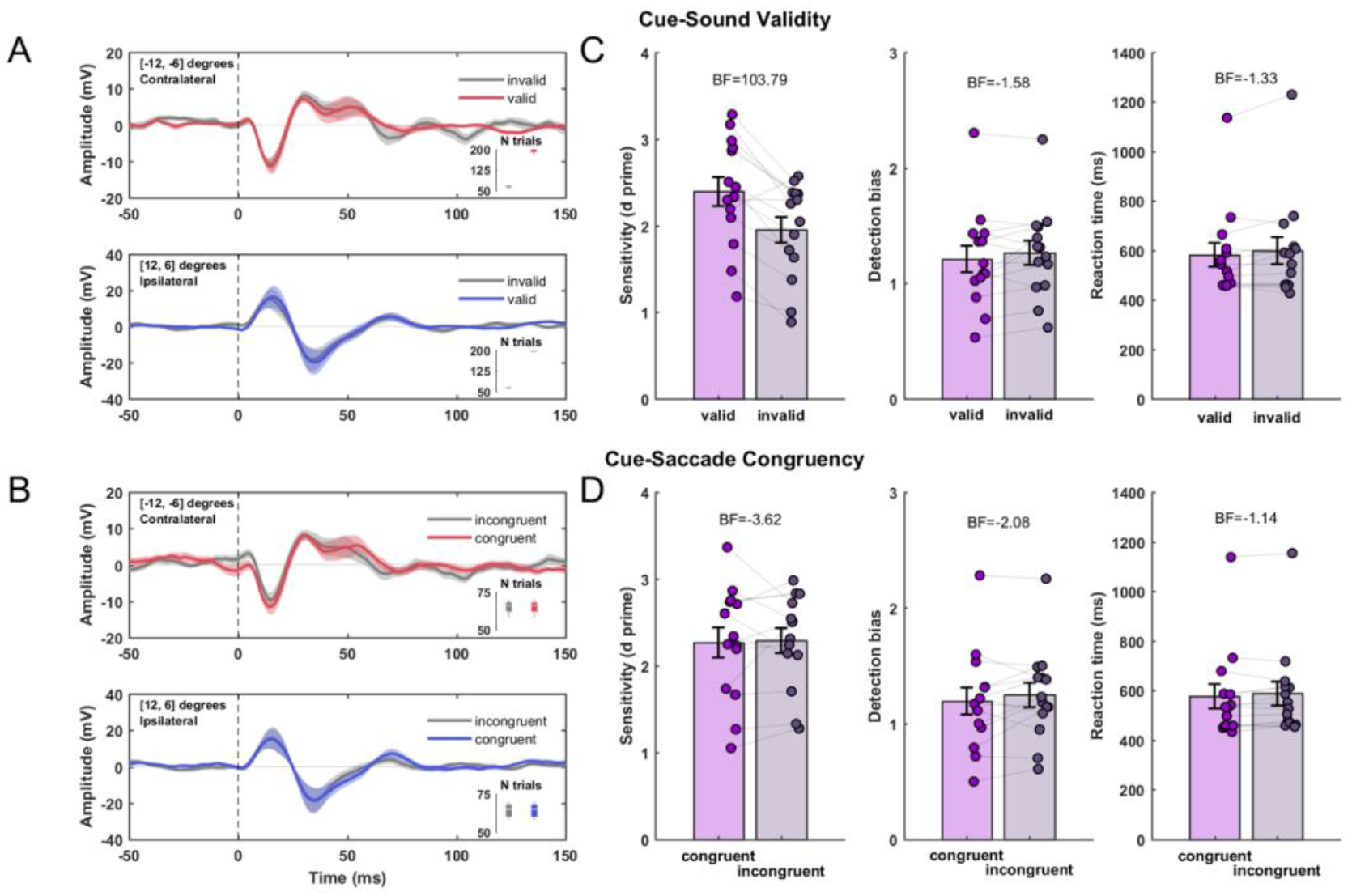
Guiding covert attention affects sound detection but not EMREOs. We probed the influence of cue-sound validity on EMREOs (**A**) and of the cue-sound congruency (**B)**; i.e. whether cues were pointing towards the same location as the saccade was directed at. The left column shows the mean EMREOs for trials divided within a factor, separate for contralateral ([-12, -6] degrees; top, red) and ipsilateral saccades ([6, 12] degrees; bottom, blue). Lines indicate the mean and shaded areas indicate the SEM across participants. (**C)** Behavioral performance analyzed for an influence of cue-sound validity and (**D**) cue-saccade congruency. Dots represent the detection sensitivity (left), detection bias (middle) and median reaction time (right) for individual participants.

Analyzing behavioral performance we found that participants exhibited higher sensitivity in trials with a valid compared to an invalid cue (Fig. 5, **C**, T(13)=-4.84, p_FDR_=0.002, BF=103.79, cohen’s D=1.3), while bias (**C**, T(13)=1.44, p_FDR_=0.26, BF=-1.58, cohen’s D=0.39) and reaction times did not differ between valid and invalid trials (**C**, T(13)=1.59, p_FDR_=0.26, BF=-1.33, cohen’s D=0.43). However, the congruency between saccade direction and sound location did not influence sensitivity (**D**, T(13)=0.23, p_FDR_=0.82, BF=-3.62, cohen’s D=0.06), bias (T(13)=1.17, p_FDR_=0.32, BF=-2.08, cohen’s D=0.31) or reaction times (T(13)=1.72, p_FDR_=0.26, BF=-1.14, cohen’s D=0.46). Hence while a valid spatial cue enhanced perceptual sensitivity this cue did not affect the EMREOs.

## 4. Discussion

We constantly scan our environment by moving our eyes. This brings new objects into our visual focus but also changes the alignment between visual signals received in retinal coordinates and acoustic information received relative to the head. Our brain therefore needs to actively align the sensory information across modalities to facilitate the encoding of congruent object-related information. The eye-movement related eardrum oscillations (EMREOs) may be a signature of such an active hearing process. However, the precise neurobiological substrate generating the EMREO and the functional implications of these for hearing remain unclear (Gruters et al., 2018; Murphy et al., 2020; Lovich et al., 2022; King et al., 2023). Our data corroborate that the eardrum moves consistently and specifically in relation to saccadic eye movements. Moreover, this directional-selective vibration of the eardrum seems to be specific to overt shifts of attention via eye movements, as our data show that covertly shifting spatial attention does not modulate the EMREO. Importantly, we also find that individual deflections of the eardrum as reflected in the EMREO do not affect the detection of near-threshold sounds. The behavioral data showed neither a difference in behavioral indices when targets were presented prior to, during or after a saccade, nor did fluctuations in the EMREO relate to detection performance. While this does not rule out that the deflection of the eardrum during an EMREO affects listening performance in more complex acoustic tasks, it shows that the most basic form of sound detection is not shaped by saccadic behavior or its signature in the ear.

### The EMREO is a reliable signal reflecting saccadic eye movements

Following previous work we recorded and aligned ear-derived microphone signals to visually guided horizontal and vertical saccadic eye movements. The EMREO time courses reported here are generally similar to those described in previous studies (Gruters et al., 2018; Murphy et al., 2020; Lovich et al., 2022). That is, the early deflections are stronger for larger saccades and their phase inverts with saccade direction. Overall the time courses in our data are somewhat slower compared to previous reports, although the overall duration of the EMREO was largely comparable. For example, the first peak occurred at around 15 ms after saccade onset in the present data and within the first 10 ms in previous work (Gruters et al., 2018), despite the hardware and data processing routines being largely identical. Importantly, both our and previous data show that the EMREO time course outlasts the saccade duration, suggesting that this signal is not a direct copy of visuo-motor commands but rather the signature of an active auditory process based on oculomotor signals.

Our data also show a considerable heterogeneity in the individual data, both in terms of relative amplitude and time course. At present it remains unclear whether this individual variability is simply measurement noise, for example arising from slightly different positions of the microphone in the ear canal, differences in ear canal diameter or whether it reflects genuine individual differences. Further studies probing the between-session consistency of the EMREO within an individual are required to address this issue.

### Sound detection is invariant to saccadic eye movements

To probe whether oculomotor behavior and the resulting EMREOs do have an influence on sound detection we systematically probed the relations of saccade parameters and EMREO amplitudes to three measures of participants’ performance. This revealed no evidence that saccade direction, distance or the relative direction of the saccade to the position of the sound was relevant for detection. Interestingly, recent findings have shown that predictable targets suppress oculomotor activity, which has been speculated to enhance perceptual performance of upcoming sensory events (Abeles et al., 2020). In contrast, we find that perisaccadic detection remains unaffected by eye movements. Additionally, we found evidence against a differential sensitivity for sounds played before, during or after saccades, which appends previous results that compared tone detection thresholds between trials with saccades and continued fixation (Harris and Lieberman, 1996). Only when we introduced a spatial cue in the paradigm did the relative timing of saccade and sound presentation matter: here participants were faster when the sound was presented later in the trial. However, in our view this probably does not reflect a relevance of the saccade itself but simply shows that the more time participants took to process the cue the faster they responded on trials where they correctly detected a target sound.

### Sound detection is invariant to EMREOs

While sound detection was not directly modulated by oculomotor behavior, residual variations in the EMREO still may influence how a sound is encoded and eventually how it is perceived. Since EMREOs reflect movements of the eardrum and sounds are transduced by movements of the eardrum, such an influence is easily conceivable. However, we did not find a difference in EMREOs between trials with correct and wrong responses, suggesting that variations in the eardrum position reflected by variations in the EMREO are not detrimental to the present detection task. Still, the absence of direct behavioral consequences of EMREOs does not speak against the idea that EMREOs could enhance or interfere with the encoding of more complex acoustic features. In fact, the perceptual task imposed here could be solved based on monaural cues only, and it could be that EMREOs shape how binaural cues relevant for spatial hearing such as interaural differences in time and intensity are encoded (Mills, 1958; Hafter et al., 1977; Smith and Price, 2014; Brown et al., 2018; Cho et al., 2023).

### EMREOs remains robust towards attentional manipulation

Given that EMREOs reflect the spatial orienting of attention by overt eye movements one could hypothesize that also covert changes in spatial attention may affect the EMREO as well. Indeed, theta rhythmic activity measured from the cochlea has been suggested to reflect cortical changes of selective attention (Köhler et al., 2021; Gehmacher et al., 2022). We here tested the attentional influence based on a traditional spatial cueing paradigm. In particular, if covert attention would similarly influence EMREOs as eye movements, the time course should show an attenuated amplitude in incongruent compared to congruent trials. However, the observed EMREOs were invariant to changes in spatial attention although participants behavior showed that they indeed exploited the cue to facilitate behavior. Our results hence suggest that auditory spatial attention is not directly mediated through EMREOs.

### Functional aspects underlying EMREOs

Since its discovery EMREOs have been considered to reflect active sensing, hence processes that adjust auditory sensitivity towards specific locations and which help to align visual and acoustic information between eye-centered and head-centered coordinates (Gruters et al., 2018). Still, the precise manner in which the eardrum is set into movement during the EMREO remains unclear. One possibility is that muscles of the middle ear, the tensor tympani and the stapedius muscle, directly drive this phenomenon. These muscles can modulate the transmission impedance of the ear, for example to protect the auditory system from very loud noises. However, they can also systematically modulate temporal transmission delays from the eardrum to the inner ear. A pivotal recent study shows that changing the tension of middle ear muscles can modulate the transmission delays from the eardrum onto the inner ear by about 150 μs (Cho et al., 2023). Interaural time delays provide important cues for spatial hearing (Wightman and Kistler, 1992; Smith and Price, 2014; Brown et al., 2018) and the reported delay corresponds to changes in ±15 degrees azimuth. In principle it is possible that the EMREO shapes the relative transduction of sounds in both ears so that they appear localized differently than they physically are. However, this specific hypothesis still needs to be tested and future work needs to specifically explore sounds defined by pure inter-aural timing or level differences and probe the precise localization of these.

Besides the precise generation of the EMREO in the ear, it also remains unclear where the signals commanding this originate from. One possibility is that the EMREO is generated directly based on a motor-efference copy from oculomotor control structures. However, there are two lines of evidence to suggest otherwise. First, the EMREO can clearly outlast saccadic eye movements by several tens of milliseconds making a direct oculomotor signal unlikely. And second, we found that most of the variability in the EMREO could be explained by the general direction and amplitude of the saccade (i.e. the experimental saccade condition) but not by residual trial-by-trial variations in their starting or end points. This suggests the generation of the EMREO is not directly driven by a one-to-one copy of the veridical eye information but rather by information about the planned upcoming saccade. Still, further work is required to explore whether motor planning reflecting the intention to move spatial attention or the actual execution of this are eventually inducing the EMREO.

## Credit author statement

Conceptualization: F.B., C.K.

Project administration: F.B., C.K.

Funding acquisition: C.K.

Methodology: F.B., C.K.

Software: F.B., C.K.

Formal Analysis: F.B.

Investigation: F.B.

Supervision: C.K.

Writing Original Draft: F.B.

Writing Review & Editing: F.B., C.K.

## Data and code availability

Data and code used in this study are publicly available on the Data Server of the University of Bielefeld (https://gitlab.ub.uni-bielefeld.de/felix.broehl/emreo_detection).

